# Reflective mirror-based line-scan adaptive optics OCT

**DOI:** 10.1101/2021.07.07.451545

**Authors:** Vimal Prabhu Pandiyan, Xiaoyun Jiang, James A. Kuchenbecker, Ramkumar Sabesan

**Affiliations:** Department of Ophthalmology, University of Washington School of Medicine, Seattle, WA, 98109, USA

## Abstract

Line-scan OCT, incorporated with adaptive optics (AO), offers high resolution, speed and sensitivity for imaging retinal structure and function *in vivo*. Here, we introduce its implementation with reflective mirror-based afocal telescopes, optimized for imaging light-induced retinal activity (optoretinography) and weak retinal reflections at the cellular scale. A non-planar optical design was followed based on previous recommendations with key differences specific to a line-scan geometry. The three beam paths fundamental to an OCT system – illumination/sample, detection, and reference – were modeled in Zemax optical design software to yield theoretically diffraction-limited performance over a 2.2 deg. field-of-view and 1.5 D vergence range at the eye’s pupil. The performance for imaging retinal structure was exemplified by cellular-scale visualization of retinal ganglion cells, macrophages, foveal cones, and rods in human observers. The performance for functional imaging was exemplified by resolving the light-evoked optical changes in foveal cone photoreceptors where the spatial resolution was sufficient for cone spectral classification at an eccentricity 0.3 deg. from the foveal center. This enabled the first *in vivo* demonstration of reduced S-cone (short-wavelength cone) density in the human foveola, thus far observed only in *ex vivo* histological preparations. Together, the feasibility for high resolution imaging of retinal structure and function demonstrated here holds significant potential for basic science and translational applications.

## 1. Introduction

Optical coherence tomography (OCT) offers 3D cross-sectional visualization of the retina [1]. Standard clinical OCT has high axial resolution (< 5 µm) but relatively low transverse resolution (∼15 µm). The former is limited by the coherence length of the OCT illumination, while the latter is limited by the eye’s optics and diffraction [2]. Incorporation of adaptive optics (AO) to OCT has allowed imaging the retina on a cellular scale in 3D [3-6]. Specifically, AO-OCT has been used to image different cell types and retinal layers, including the internal limiting membrane (ILM) [7-9], nerve fiber layer (NFL) [9-11], retinal ganglion cell layer (RGC) [8, 9], cone photoreceptors [11-16], rod photoreceptors [17-20], retinal pigment epithelium (RPE) [9, 19, 20], and choriocapillaris [20, 21].

Besides structure and morphometry, variants of AO-OCT have also been employed for optoretinography, or imaging the light-evoked retinal activity. This is undertaken by measuring the retinally backscattered optical phase in response to a light stimulus [16, 22-27]. The application of non-AO OCT and AOSLO to measure changes in optical intensity in response to light has also been described [28-36]. Together, these offer a sensitive and non-invasive *in vivo* assay of retinal function with profound implications for both basic and clinical science. The speed of acquisition for such measurements are universally important in order to reveal fast light-induced retinal events, especially in the presence of eye movements. Coupled with AO via hardware or computational (or software) implementations enables localizing the elicited activity to individual cells. Such implementations have ranged across a continuum of confocality and speed. On the one hand, full-field OCT enables high speed at the cost of confocality [26, 27], while on the other, point-scan swept source and spectral domain AO-OCT enable excellent resolution & contrast, at the cost of speed [22, 24, 25].

Line-scan AO-OCT affords a tradeoff between these extremes by offering partial confocality across one dimension together with high cross-sectional B-scan acquisition speeds. AO-OCT implementations in line-scan configuration have employed both digital and hardware correction, revealing retinal structure and function at high resolution. With AO, Zhang et al. [6] made the first observations of the interface between the inner and outer segments of individual cones in retinal B-scans, though the low B-scan rate of 500 Hz impeded volumetric imaging. Pandiyan et al. [16] presented the first volumetric, high-speed line-scan spectral domain OCT with hardware AO, aimed at phase-resolved acquisition of light-induced retinal activity or optoretinography.

AO ophthalmoscopes, in general, use afocal telescopes to relay light and optically conjugate pupil planes to allow for placement of scanners, deformable mirrors, and other static optical corrections like sphero-cylindrical lenses. In line-scan AO-OCT, a cylindrical lens generates a linear illumination at the entrance pupil of the system, that is then optically conjugated with a 1-D galvo scanner, a deformable mirror and the eye’s pupil using afocal telescopes. In Pandiyan et al., [16] achromatic lens-based telescopes were used for the ease of alignment and an abundant commercial availability of effective focal lengths. However, lens back-reflections are a common issue that are removed by tipping and tilting the lenses, which in turn leads to optical aberrations. The aberrations in the retinal plane are correctable with the deformable mirror, but pupil plane aberrations persist. Spherical mirror based afocal telescopes when arranged in a non-planar arrangement [37, 38] help alleviate the issues posed by afocal lens-based telescopes and eliminate back-reflections. Thus, majority of AO ophthalmoscopes use reflective optics in portions of the optical path where light traverses the optical components in double-pass. This arrangement helps improve signal efficiency by reducing losses, and is especially advantageous for removing spurious reflections in the wavefront sensor. However, a linear polarizer and a quarter-wave plate used in combination can effectively reduce lens back-reflections in a lens-based AO-OCT system [20, 39].

Here, we introduce the design and implementation of a reflective mirror-based line-scan AO-OCT. The basic principle parallels that put forth for point-scan AOSLO, with modifications aimed at optimized performance in a line-scan illumination and collection geometry for AO-OCT. The performance was evaluated by imaging retinal structure and light-evoked cone activity in human volunteers. Emphasis was kept on testing the limits of visualizing retinal structures with weak reflections (e.g.: retinal ganglion cells) and those close to the resolution limit of the eye (e.g.: foveal cones, rods). The limits for assessing the light-evoked functional response at cellular-scale spatial resolution was evaluated by performing optoretinography of the minute cone photoreceptors in the foveola.

## 2. Methods

### 2.1 Optical design and layout of the reflective line-scan adaptive optics retinal camera

Broadly, the design process consisted of the following steps. First, the specifications for the optical instrument were chosen, particularly the desired a) lateral and axial resolution, b) axial depth, and c) field-of-view (FOV) and d) vergence range, over which optical performance is diffraction-limited. Next, the optical components – source, detector, grating, spherical mirror and lens focal lengths – were chosen to meet the desired resolution (lateral, axial, and spectral) and FOV. Constraints on these choices were made based on the commercial, off-the-shelf availability of optical components and the size of the optical table (4 ft x 8 ft). Next, optical design software Zemax (Zemax LLC, Kirkland, WA, USA) was used to optimize the optical design in the sample, reference, and detection channels to achieve the desired specifications above. The candidate optical design was then exported to computer-aided design (CAD) software SolidWorks (Dassault Systèmes SolidWorks Corporation, Waltham, MA, USA) where the optomechanical mounts were added to assess mechanical restrictions and beam vignetting. The optical and mechanical considerations were iteratively optimized using the combination of SolidWorks and Zemax to arrive at the final optomechanical design consisting of the xyz-coordinates of each optical component. Fig.1 shows the optical layout of the mirror-based line-scan AO camera, not drawn to scale. The specifications of the system are listed in Table 1. The focal lengths of the spherical mirrors and lenses are listed in Table 2. Below, we detail the design considerations and optical design of the three channels - illumination, detection, and OCT reference path separately.

**Fig. 1.**
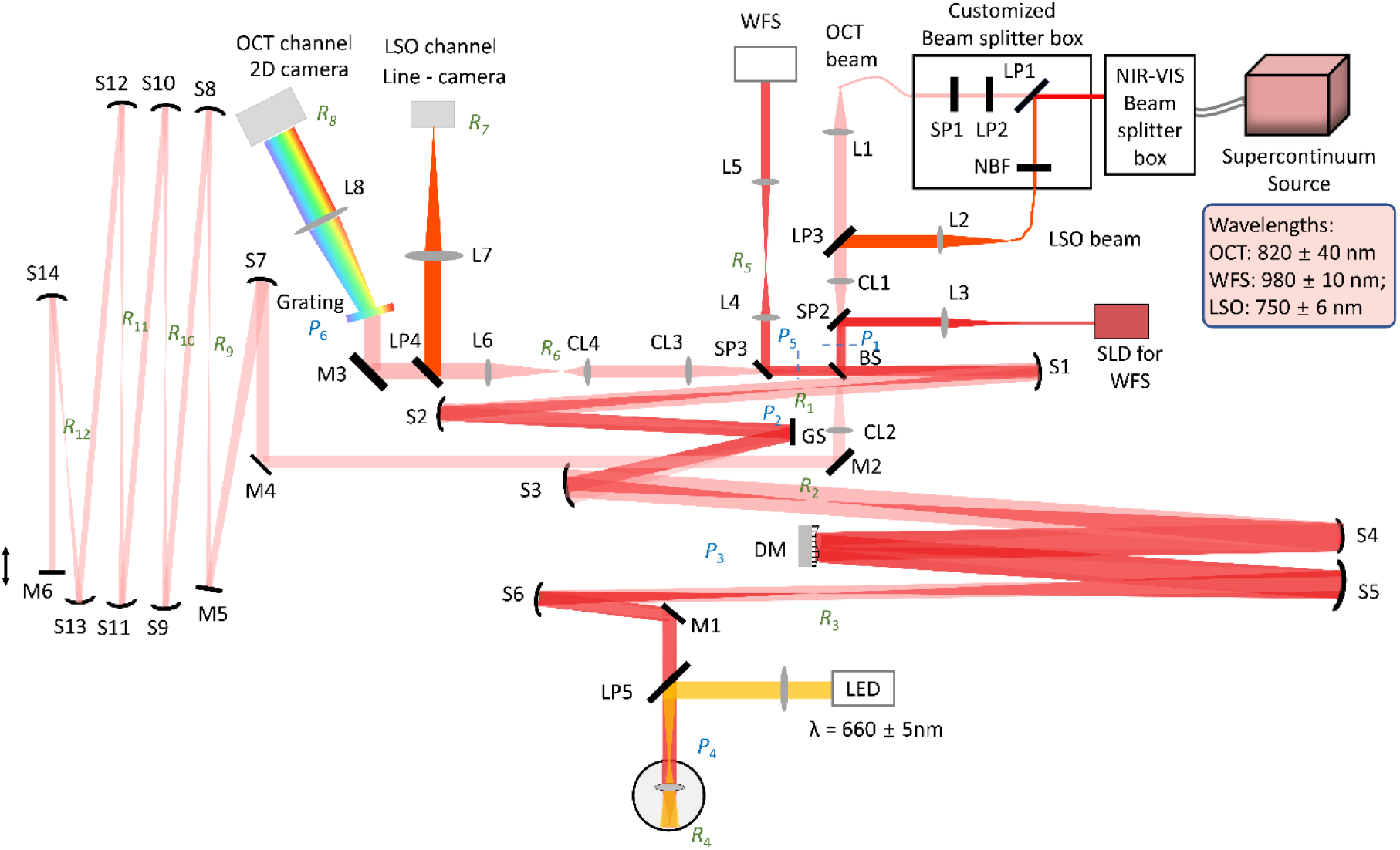
Optical layout of the LS-AOOCT/LSO. L: lens; S: Spherical mirror; M: Flat mirror; BS: Beam splitter; LP: Long-pass filter; SP: Short-pass filter; NBF: Narrowband filter; GS: Galvo scanner; DM: Deformable mirror; WFS: Wavefront sensor; P: Pupil plane; R: Retinal Plane

**Table 1.**
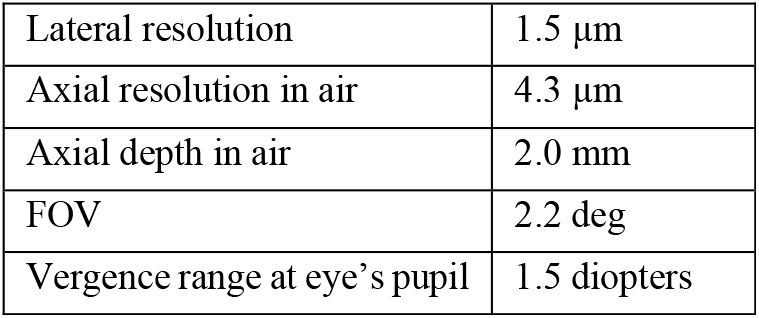
Specification of the system.

|  |  |
| --- | --- |
| Lateral resolution | 1.5 $\mu\text{m}$ |
| Axial resolution in air | 4.3 $\mu\text{m}$ |
| Axial depth in air | 2.0 mm |
| FOV | 2.2 deg |
| Vergence range at eye's pupil | 1.5 diopters |

**Table 2.** List of spherical mirror focal lengths and detection arm focal lengths.

| Optical element | Focal length(mm) | Optical element | Focal length(mm) |
| --- | --- | --- | --- |
| CL1 | 75 | S10 | 500 |
| CL2 | 100 | S11 | 500 |
| CL3 | 250 | S12 | 500 |
| CL4 | 75 | S13 | 500 |
| S1 | 500 | S14 | 250 |
| S2 | 750 | L1 | 50 |
| S3 | 500 | L2 | 40 |
| S4 | 1000 | L3 | 30 |
| S5 | 1000 | L4 | 150 |
| S6 | 500 | L5 | 200 |
| S7 | 750 | L6 | 300 |
| S8 | 500 | L7 | 300 |
| S9 | 500 | L8 | 350 |

#### 2.1.1. Illumination Path

A supercontinuum light source (SuperK EXR-9OCT, NKT Photonics, A/S, Birkerød, Denmark) operating at a repetition rate of 320 MHz was chosen based on its ability to provide sufficient output spectral power density, flexibility in choice of central wavelength and spectral bandwidth. This source has been characterized previously for use in OCT [40]. A passive spectral splitter (SuperK split spectral splitter, NKT Photonics, A/S, Birkerød, Denmark) combined with a custom in-house spectral splitter was used to derive two imaging wavelength bands: 820 ± 40 nm for OCT and 750±6 nm for line-scan ophthalmoscope (LSO). The in-house spectral splitter consisted of a network of short (SP1) & long-pass (LP1 and LP2) filters, and a narrowband filter (NBF). A 980±10 nm superluminescent diode (SLD) (IPSDD0906, Inphenix, USA) was used for wavefront sensing. The OCT and LSO beams were combined by a long-pass filter (LP3) prior to a cylindrical lens (CL1). Both formed a linear illumination profile at the entrance pupil (P_1_), while the wavefront sensing beam joined the main illumination path via a short-pass filter (SP2) after the cylindrical lens (CL1) and formed a circular beam profile at the entrance pupil (P_1_). A beamsplitter (30:70, reflectance: transmission ratio) separated the OCT sample and reference arms. Three spherical mirror-based telescopes were arranged in a non-planar off-axis configuration to optically conjugate the entrance pupil to the eye’s pupil (P_4_) via two intermediate pupil planes – dedicated to a 1-dimensional galvo scanner (GS, P_2_) and the deformable mirror (DM, P_3_) respectively.

The optical performance was optimized based on the general method described by Dubra et al. [41]. We have elaborated previously the manner in which the illumination and detection beam profiles differ in line-scan geometry. For completeness here, note the differences in fig. 2 in the cross-sectional view for the illumination and detections paths at several pupil and retinal planes. A cylindrical lens creates a linear beam profile in every pupil and retinal plane with their orientation rotated by 90°. At each plane, the beam is focused along one dimension. In detection, most notably, the linear field illuminating the retina loses its phase-front and gives rise to a set of beamlets with near-spherical wavefronts when backscattered. The backscattered light overfills the eye’s pupil and results in spherical beam profiles at the pupil planes in detection. Given this difference in beam propagation in illumination versus detection, their performance was optimized and assessed separately.

**Fig. 2.**
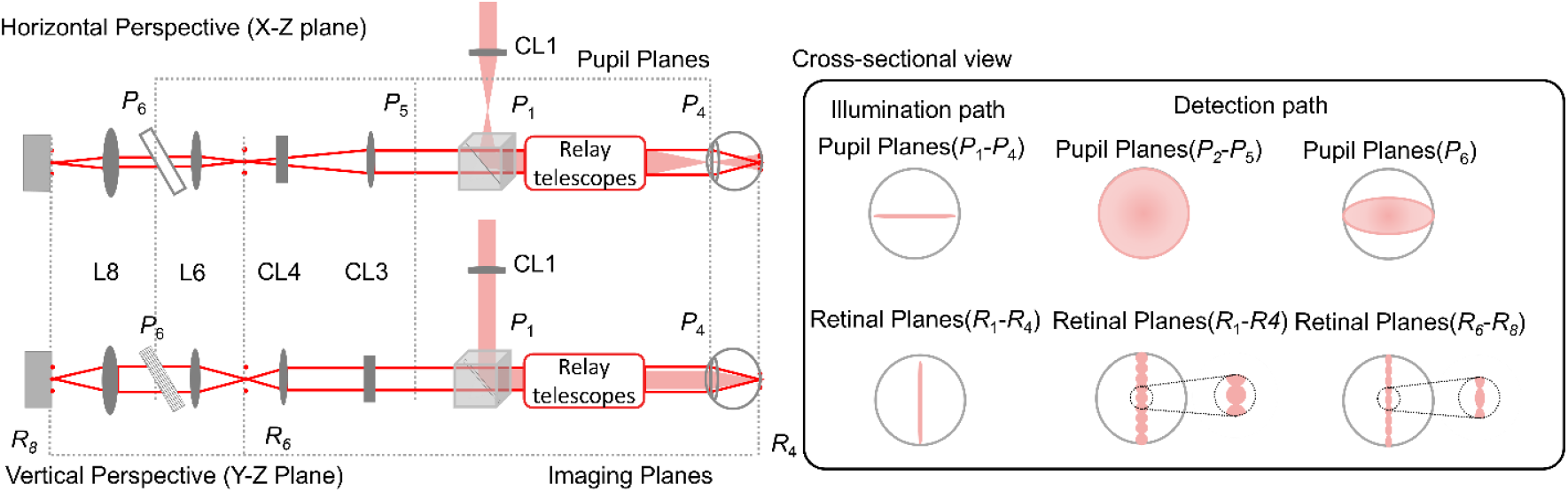
The horizontal and vertical perspective optical path for both illumination (red shaded) and detection (red solid line) are shown with the same naming convention as fig 1. In the illumination path, the beam profiles at both pupil and retinal planes are linear with Gaussian distribution along the line dimension. In the detection path, note the beam profile differences at pupil and retinal planes before and after the anamorphic configuration (CL3 and CL4). Before the anamorphic configuration, the beam profiles are circular at the pupil planes(*P*_2_-*P*_5_), while afterwards at the grating (*P*_6_), the beam profile is elliptical. In detection, a line profile of circular and elliptical beamlets are made at the retinal planes(*R*_1_-*R*_4_) and (*R*_6_-*R*_8_), before and after the anamorphic configuration respectively.

Fig. 3a and 3b show the illumination spot patterns after optimization at the final eye’s pupil and retinal planes respectively along the dimension where the beam is focused. Diffraction-limited performance was achieved over the 2.2 deg. FOV and 1.5 D vergence at the eye’s pupil. This vergence range covers the longitudinal chromatic aberration between 750 to 1000nm spectral range (0.4 D), and the optical power (0.9 D) needed to section the retina.

**Fig. 3.**
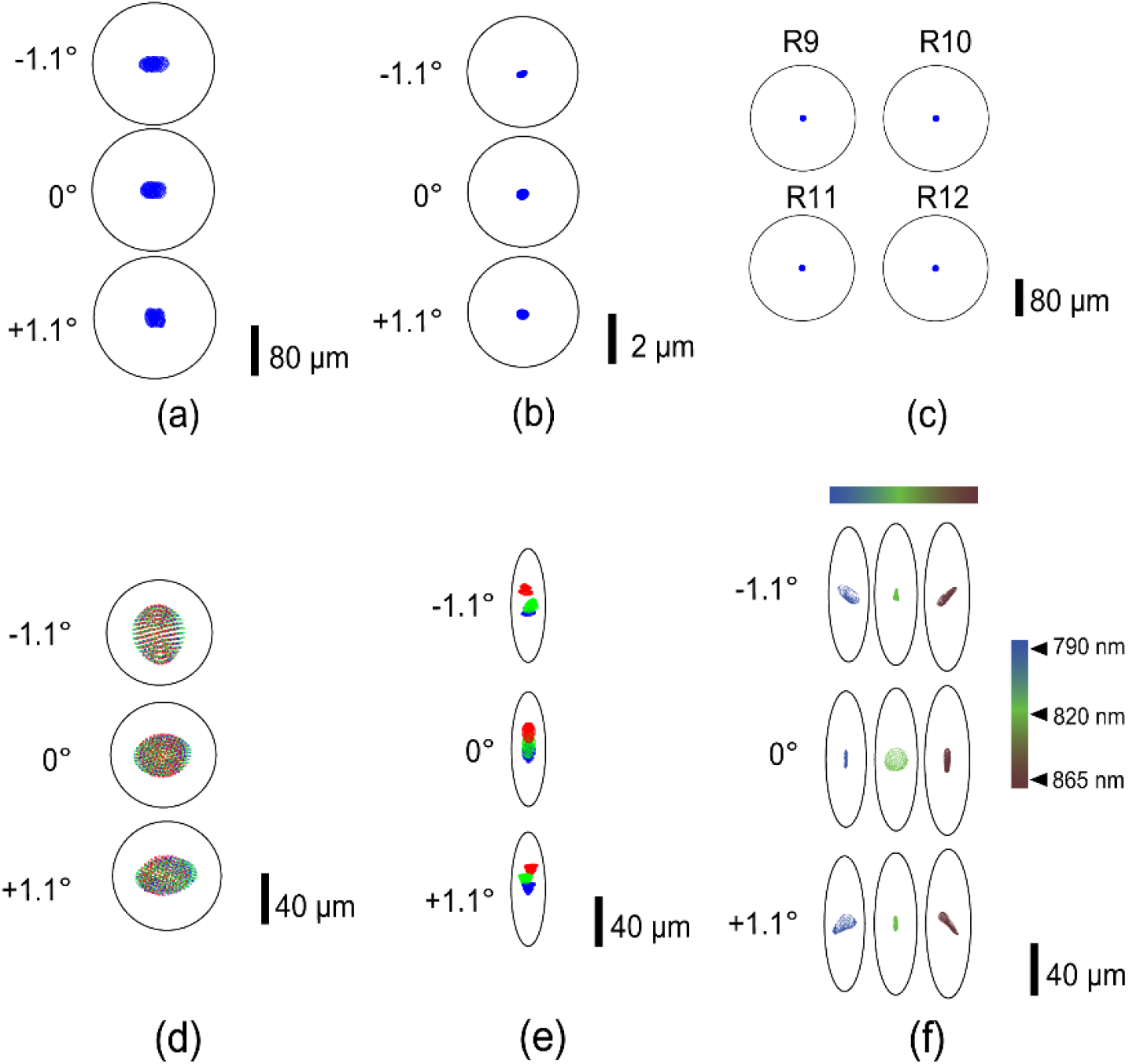
Spot diagram for (a) illumination path pupil plane (P4) (b) illumination path retinal plane (R4), (c) reference arm retinal plane (R9-R12) (d) Detection path pupil plane (P5) (e) Detection path retinal plane (R6) (f) Detection path retinal plane (R8) (d,e) green, blue and red (−1.1°,0°,1.1°) represents three field points along the line dimension. In (f) the color bar and the corresponding spot patterns indicate performance across the spectral bandwidth of OCT source illumination

#### 2.1.2. Detection path

The detection path was modeled in Zemax starting with three field points (−1.1°, 0° and 1.1°) at the eye’s retinal plane, to mimic the array of beamlets distributed across the line-scan and backscattered from the retina. A virtual scanner plane was created at the eye’s pupil plane that had an equal and opposite scan angle as the galvo-scanner, such that retinal planes R2 and R3 had square imaging fields in the detection path, and the three fields along the scan dimension were descanned after the galvo-scanner to yield a linear field at retinal plane R1. The three fields created a circular profile at each pupil plane until the exit pupil P5. The established Zemax model remained same with respect to the xyz coordinates and tip/tilt angles in both illumination and detection. The optimized Zemax performance at the exit pupil P5 and retinal plane R6 along the scan field are shown in fig. 3d and 3e. The different colors in the figure represents the field points (−1.1°, 0° and 1.1°) along the line dimension.

An anamorphic configuration consisting of two-cylinder lens achromats (CL3 and CL4 in Fig.1) was applied to optimize the spectral and spatial resolution, and to improve light collection efficiency. The principle behind its operation is elaborated in Pandiyan et al. [16]. This configuration provided an asymmetric beam magnification such that following pupil (P6) and retinal (R6 – R8) planes were scaled by the ratio of the focal lengths of CL3 and CL4. Accordingly, the diffraction limited airy disk is transformed in an *‘airy ellipse’* in fig. 3(e,f). A 600 line-pairs per mm linear transmission grating (WP-600/840-35×45, Wasatch Photonics, USA) was used to build a customized spectrometer. The Zemax model in the detection arm was optimized based on the optical performance at the final OCT imaging plane (2D camera) for each field along the line dimension and wavelength configuration to yield diffraction-limited spot patterns, as shown in Fig 3(f). For simplicity, only the 0° field point along the scan dimension is shown here.

#### 2.1.3. OCT reference arm

The aberrations introduced by the off-axis use of spherical mirrors, specifically in afocal telescopes scale inversely with the mirror focal lengths. Hence, we chose focal lengths from 250 – 750 mm, yielding an overall double-pass sample arm length of 8.5 meters. An identical reference arm optical path length was thus needed to match the sample arm. In addition, the cylindrical phase front and diffraction needed to be minimized over the long travel for optimal interference. Three spherical mirror-based telescopes were used to relay the beam, and simultaneously reduce diffraction and eliminate dispersion. The spot diagrams were easily optimized by adjusting the tip-tilt angles of the spherical mirrors to provide diffraction limited performance at each imaging plane in the reference arm, as shown in fig.3(c).

### 2.2 Imaging protocol – acquisition and processing

Two cyclopleged subjects free of retinal disease were recruited for the study after an informed consent explaining the nature and possible consequences of the study. The research was approved by the University of Washington institutional review board and the experiment was performed in accordance with the tenets of the Declaration of Helsinki.

Imaging was conducted at several eccentricities from the foveal center to 10 deg. temporal. Focus was set to either the outer or inner retina, assisted by a real-time video stream in the LSO channel. The focusing lens (L7) and line-scan camera (Basler, sprint, spL2048-70km pixel size 10 µm) for the LSO were integrated in the detection path by using either a long-pass filter or a removable mirror in the optical path, and optically conjugated with the OCT camera.

A Shack-Hartmann wavefront sensor (lenslet pitch: 200 µm, lenslet focal length: 9 mm, CCD camera pixel size: 6.45 µm) was placed in reflection path of a short-pass filter (SP3) and used to measure the aberrations for closed-loop AO operation. An adjustable iris was placed at R5 to minimize corneal backreflections. In illumination and detection, the wavefront sensor beam was restricted by the deformable mirror (DM97-15, Alpao, France) aperture, equal to 6.75mm at the eye’s pupil. At the wavefront sensor camera, this beam was demagnified to cover 6 mm.

Imaging performance in the line-scan AO-OCT was evaluated under three different conditions where the acquisition protocol varied as follows:

1. For imaging the foveal cones, 1.5 deg temporal eccentricity and foveal cone ORG, the B-scan rate was 12000 scans/sec. Each volume contained 600 B-scans, and each video contained 50 volumes. Six recordings were taken. For the ORG experiments, subjects were dark adapted for 3 minutes and data were recorded with 660 nm, 20ms light flash delivered after the 10^th^ volume. The OCT phase analysis post-acquisition followed from a previous article [16].
2. For imaging the RGC layer, the B-scan rate was 6000 scans/sec and volume rate was 10 volumes/sec. Each volume contained 600 B-scans, and each video contained 20 volumes. Ten recordings were taken at 3-minute intervals.
3. For imaging the RPE layer, the B-scan rate was 12000 scans/sec and volume rate was 20 volumes/sec. Each volume contained 600 B-scans, and each video contained 50 volumes. Six to ten recordings were taken at 6-minute intervals.

In all cases, typical processing steps – background subtraction, k-space resampling, Fourier transform, image registration and segmentation – were followed, and detailed in reference [16].

## 3. Results

Here, we show exemplary images from the instrument at different eccentricities, and retinal layer foci. Next, we show an example application for imaging the light-evoked activity in individual cone photoreceptors near the foveal center.

### 3.1 Structure imaging

Fig. 4a shows an average AO-LSO image in logarithmic scale. In this image, cones are resolved near the foveola, but the foveal center eluded visualization. Fig. 4b shows maximum intensity projection of 5 pixels centered around the inner-outer segment junction (ISOS) and the cone outer segment tips (COST) at the foveola recorded at the speed of 20 volumes/sec (vol/s) where the focus was set to the outer retina. Fig. 4c shows a magnified view of fig. 4b. Cone photoreceptors are resolved readily up to 0.3° from the foveal center. Cones still nearer to the foveola were beyond the resolution limit, likely due to insufficient digital resolution or optical resolution. Nyquist sampling (0.15 arcmin per pixel in retina) was kept low in favor of improved light collection per pixel. Note that this subject’s axial length - 24.33 mm - was slightly longer than the population average (24 mm; [42]). Theoretically, a shorter axial length improves lateral resolution by virtue of a correspondingly higher numerical aperture.

**Fig. 4.**
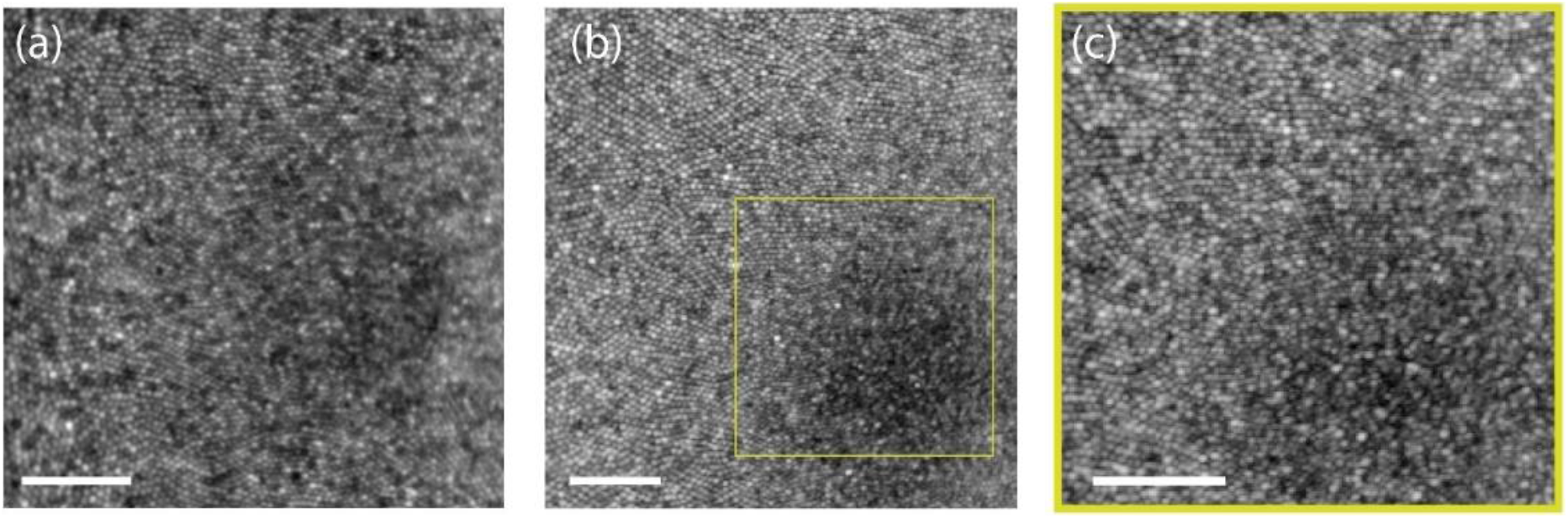
(a) Foveal LSO image at 720 nm (b) *En face* projection of ISOS and COST from the registered AO-OCT volumes. (c) Magnified view of yellow square shown in (b). All the images are shown in log scale. Scale bar: 10 arcmin

Figs. 5a and 5b show a B-scan from an averaged AO-OCT volume at 1.5° temporal eccentricity in linear and log scale recorded at 20 vol/s, where the focus was set to the outer retina. The cross-sectional image of retina reveals the cone photoreceptor ISOS and COST layers. Maximum intensity projections at the ISOS and COST reveal cone photoreceptors with adequate resolution as shown in fig. 5c and 5e. Further, at ∼6-10 µm below the ISOS reflection, a sparse subset of reflections appeared as shown in fig. 5d. These were previously suggested to be S-cones that have shorter outer segments compared to L- and M-cones [43-45].

**Fig. 5.**
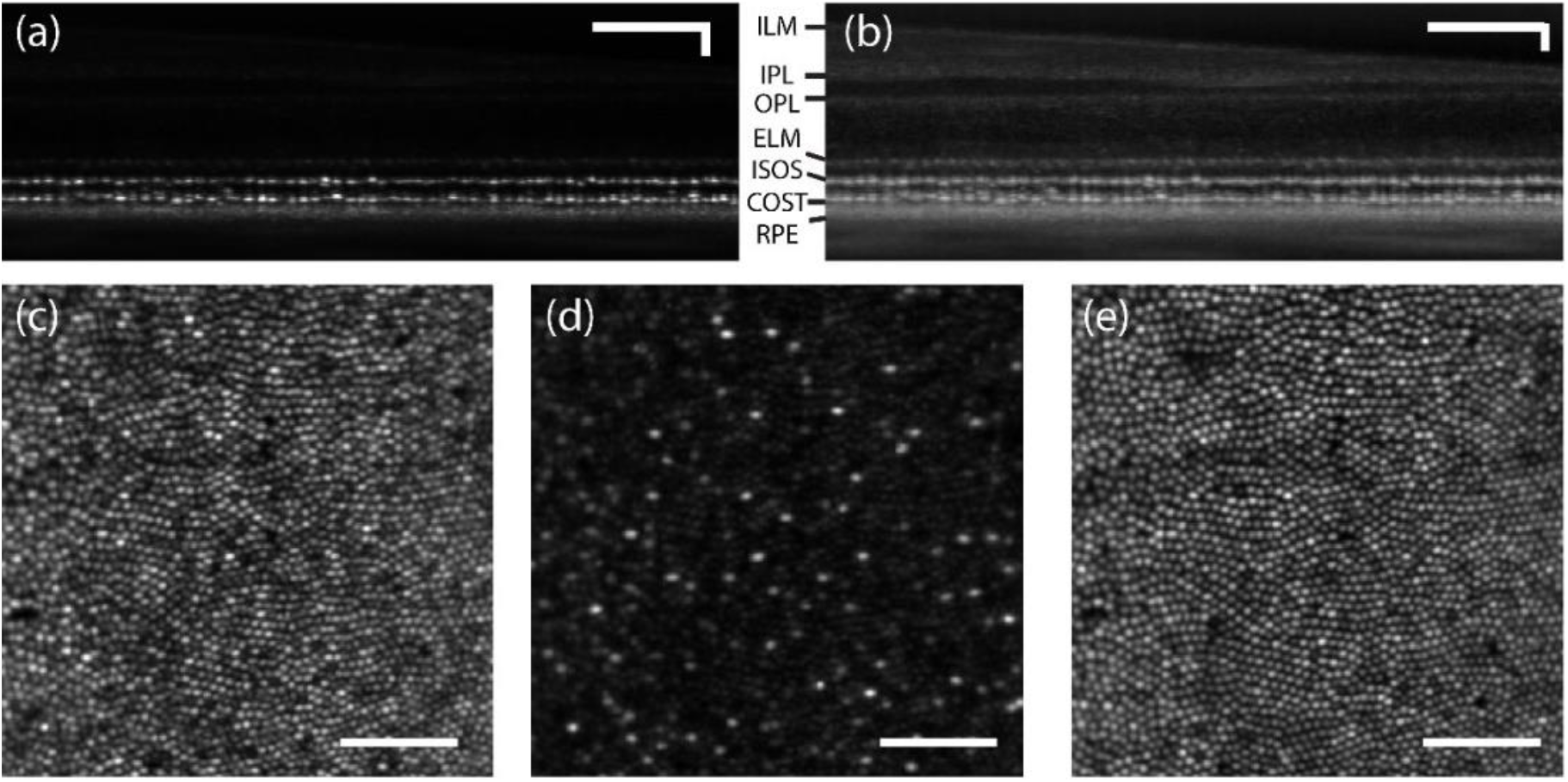
AO-OCT B-scan of retina at 1.5° temporal eccentricity in linear (a) and log (b) scale. (c) Maximum intensity projection at ISOS (d) *En face* projection 6-10 µm distal form the ISOS reflection (e) Maximum intensity projection at COST. Scale bar: 10 arcmin.

Fig. 6 shows the AO-OCT *en face* projections in linear (fig. 6(a-d)) and log scale (fig 6(e-h)) obtained at 10 deg. temporal eccentricity with the focus set to the outer retina. Here, the ISOS *en face* projections (fig. 6a, 6e) show waveguiding modes, typically observed from cone ISOS reflections beyond 4 deg eccentricity [46]. A majority resemble linearly polarized LP11 modes, however no quantitative treatment was followed here to assess the extent of mode superposition and content. In the COST *en face* images (fig. 6b,6f), relatively fewer reflections are associated with a dominant linearly polarized mode. Further distal to the COST, the rod outer segment tips (ROST) shown in the fig. 6d and 6h are visualized with cellular resolution. Here, the individual ROST encircle a darker shadow cast by the neighboring cones. Note that the ROST projection, when shown in log scale fig. 6h, accentuates a few brighter COST reflections from the background and further aid the visualization of individual rods. Further distal to the ROST, the retinal pigment epithelium (RPE) mosaic is revealed (fig. 6c, 6g), the contrast in which is suggested to be a result of organelle motility [47]. Ten videos were taken at 6-minutes intervals to capture this organelle motility. Figs. 6(i-k) show the 2D log10 power spectrum of ISOS, COST and RPE respectively. A radially averaged power spectrum (fig. 6l) shows a notable peak corresponding to COST & ISOS, appearing at a greater spatial frequency compared to the peak corresponding to the RPE sub-mosaic. The manually identified power spectrum peak at ISOS and COST matches well, equal to 8441 cones per mm^2^, which is in close agreement to a direct count equal to 8280 cones/mm^2^. Similarly, the power spectrum peak of the RPE sub-mosaic corresponds to 5549 cells/mm^2^ that is in close agreement to a direct count of 5926 cells/mm^2^. At 10 deg eccentricity, this observed density of cones relative to RPE is consistent with prior work [48].

**Fig. 6.**
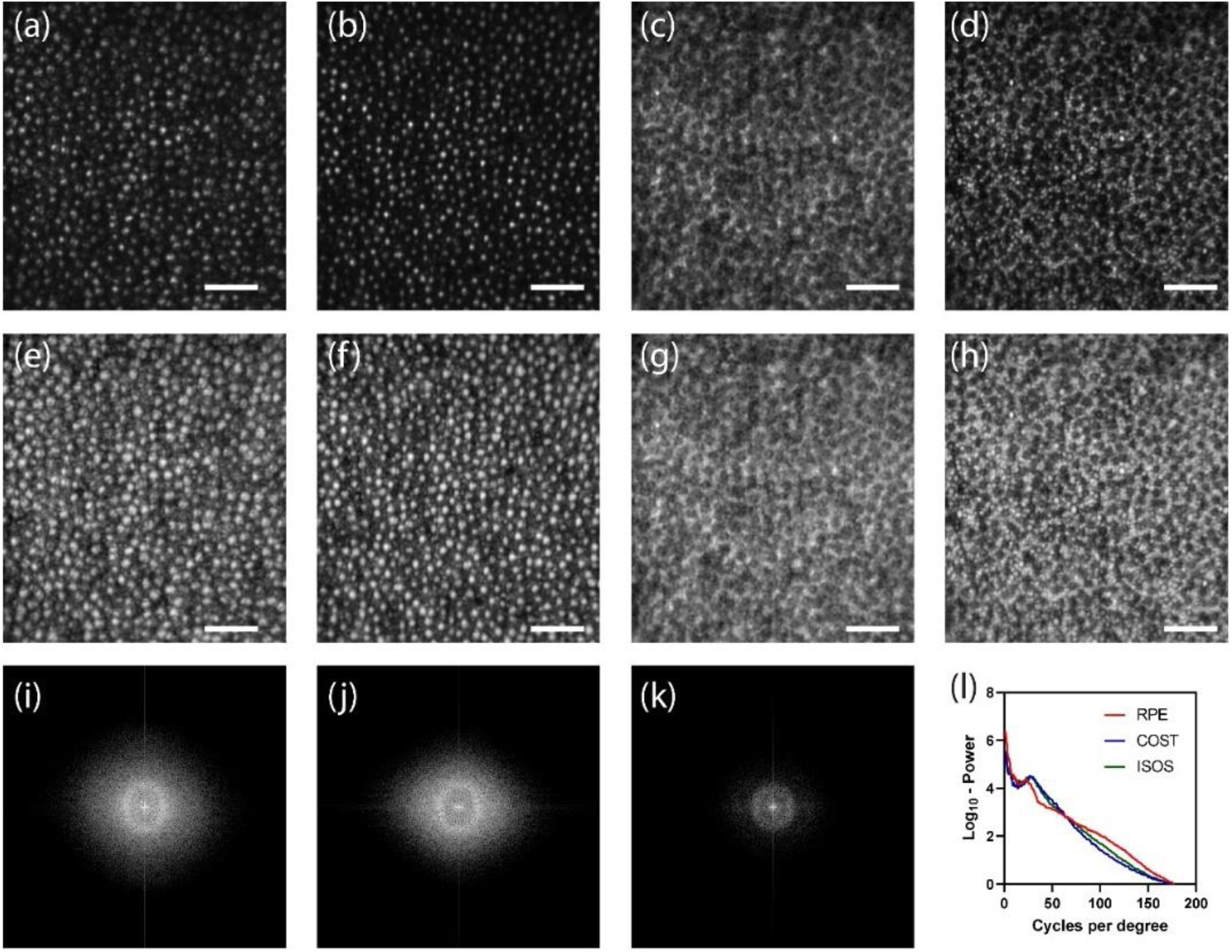
*En face* projection of ISOS (a,e), COST (b,f), RPE (c,g) and ROST(d,h) in linear (a-d) and log scale (e-h). 2D log10-power spectrum of ISOS (i), COST (j) and RPE (k). Radial average of the log10-power spectrum of ISOS, COST and RPE. Scale bar: 10 arcmin.

Fig 7 shows the AO-OCT *en face* projections obtained from the same averaged volume at 10° temporal eccentricity with the focus set in the inner retina. Ten videos were taken at 3-minute intervals to capture the organelle motility within retinal ganglion cells. Depth-resolved projections reveal several structures within the inner retina – (a) macrophages in the ILM, (b) nerve fiber bundles in the NFL, (c) retinal ganglion cells in the RGCL and (d) blood vessels in the INL. The macrophages seen at the surface of the ILM have been previously referred to as such, and no distinction can currently be made between these structures, hyalocytes, or microglia [7]. The RGCs appear in at least two spatial sizes - purported to be of midget and parasol types.

**Fig. 7.**
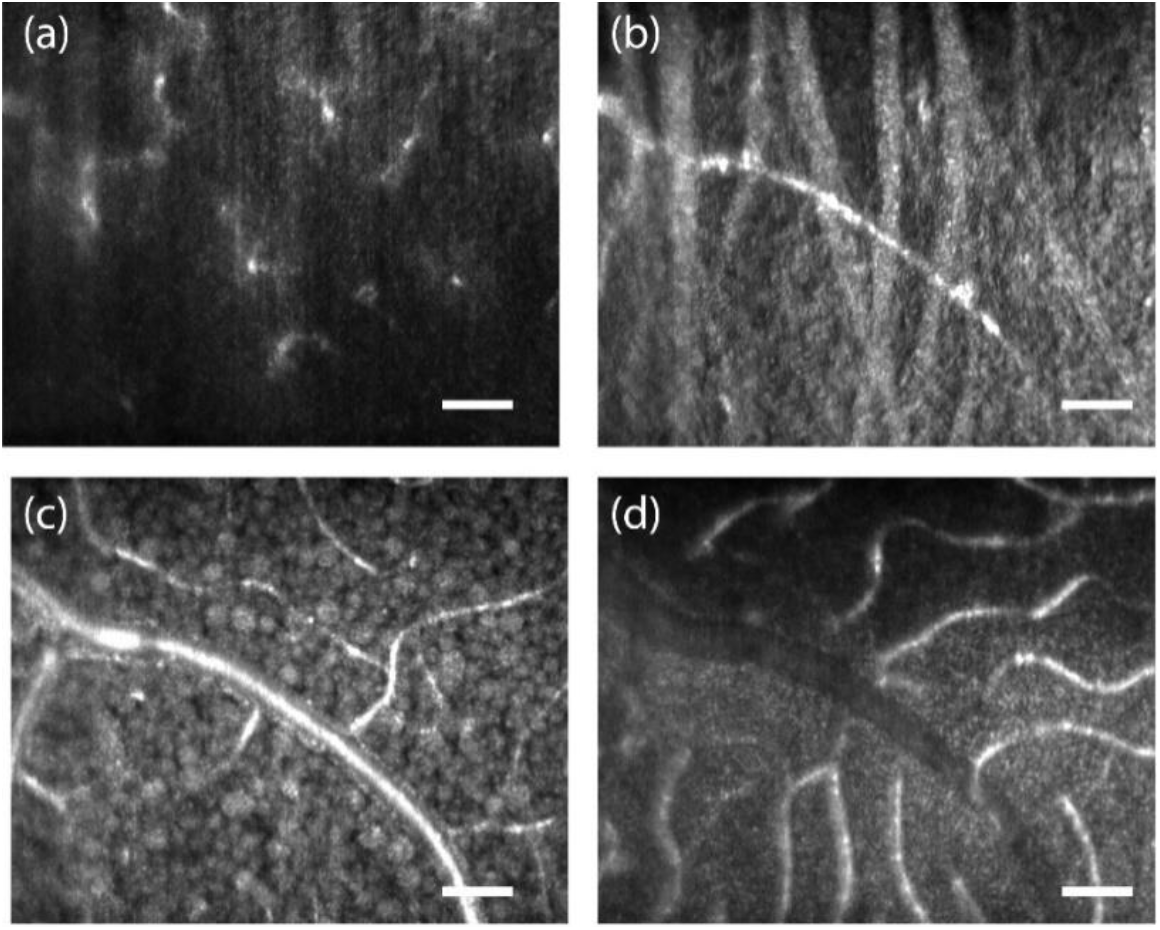
*En face* projection of inner retina (a) Inner limiting membrane showing macrophages (b) Nerve fibers (c) Retinal ganglion cells, (d) Blood vessels. Scale bar: 10 arcmin

Fig. 8 (a-c) exemplifies the feasibility of optoretinography at the center of the foveola (marked with a yellow asterisk). The demands on spatial resolution are the greatest in the foveal center, where cones are the most tightly packed and subtend ∼0.5 arc-min visual angle. Fig. 2f shows an *en face* projection from registered AO-OCT volumes at the fovea. With 660 nm stimulus, the OPL traces of single cones readily segregate into three clusters denoting L, M and S-cones, shown for two ROIs encircling the foveola (b & c). Most interestingly, the reduced density of S-cones in the foveola, evident from previous histological literature [45], is revealed here for the first time *in vivo* in human retina. The relative % of SML cones is 1.9, 26 and 72% and 0.4, 28.5 & 71.1 % in the marked ROIs respectively, calculated over 316 and 228 cones respectively. We restricted our attention only to the area(s) around the very center of the foveola where a hexagonal cone packing was readily evident. In face of the extremely tight cone packing, the resolution is still limited to corroborate the purported absence of S-cones in the foveola. Nevertheless, a substantially lower percent of S-cones (0.4 and 1.9%) is apparent at ∼20 arc-min from the foveal center, that rapidly increases to ∼6 % at 1.5 deg. eccentricity in the same eye.

**Fig. 8.**
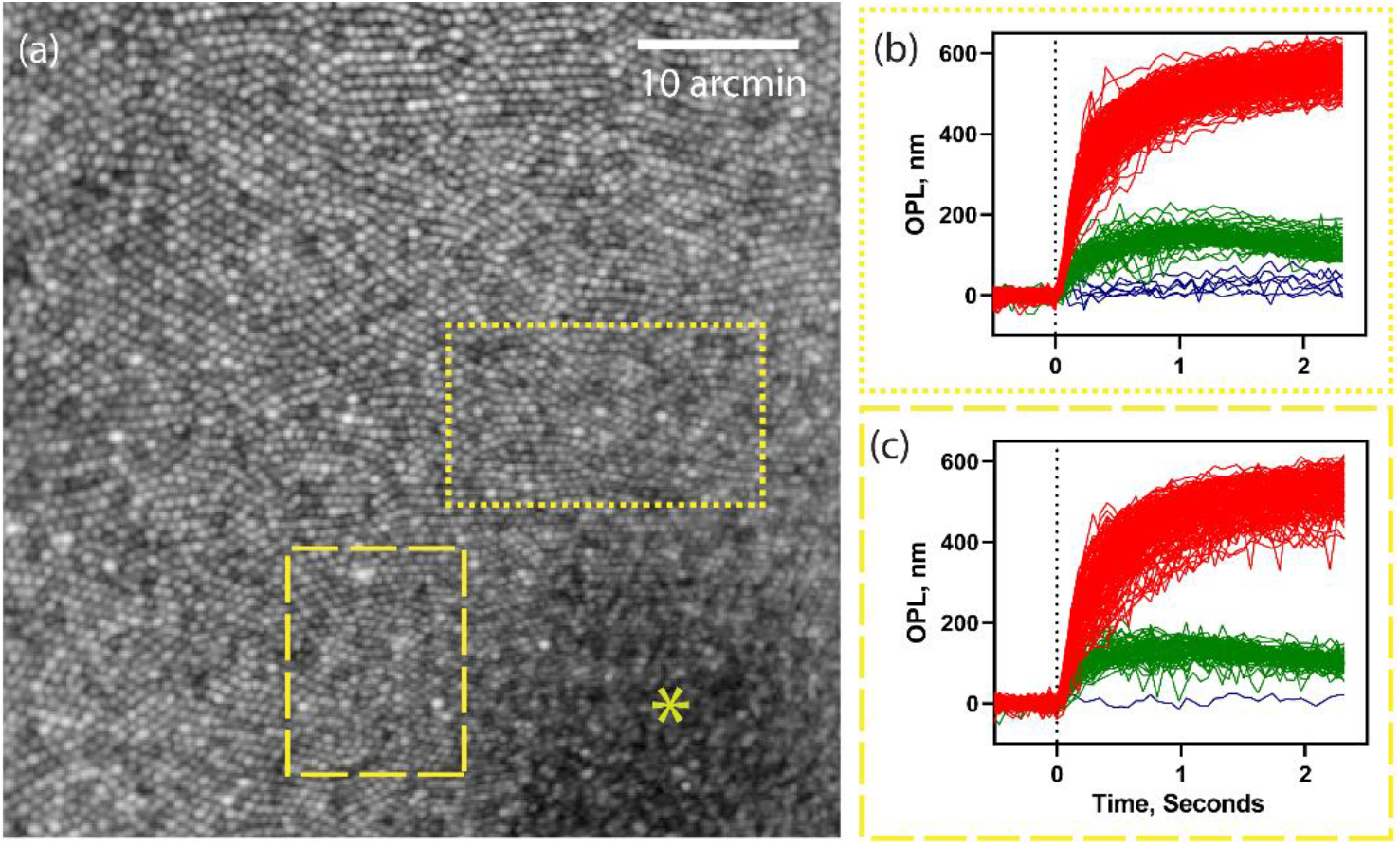
(a) *En face* projection of photoreceptors at the foveal center (b,c) Foveal cone optoretinograms corresponding to the dotted and dashed rectangular regions-of-interest in (a). Red, green, and blue color represent the response of long, middle and short wavelength cones. * - represents the foveal center.

## 4. Discussion

We implemented an AO line-scan spectral domain OCT system with reflective mirror-based afocal telescopes, aimed at cellular-scale visualization of retinal structure and function. A non-planar optical design was followed based on previous recommendations [38, 41] with key differences specific to a line-scan geometry. The three beam paths fundamental to an OCT system – illumination/sample, detection and reference – were modeled in Zemax optical design software to yield theoretically diffraction-limited performance. On the other hand, in an AO-OCT in point-scan configuration, the optical design of the sample arm and spectrometer are typically optimized to maximize performance [49]. Free-space operation and long-distance beam propagation in line-scan OCT impose extra constraints such that the OCT reference arm and the entire detection arm, including the spectrometer, additionally require optical design optimization. A supercontinuum laser was introduced for AO line-scan OCT which offered high bandwidth and spectral power density.

The resulting optical performance, as it relates to imaging retinal structure at high spatial resolution is exemplified via the following notable features. First, the resolution is sufficient to observe individual rods and foveolar cones (Fig. 4, 6 & 8 respectively) in an *en face* projection, about ∼2 µm in diameter each. Second, the weakly reflective cells in the RGCL have eluded visualization, until recently [8]. Remarkable improvements in optical design [49], acquisition speed [11] and 3D volume registration [50] together enabled imaging the morphology and distribution of RGCs. It is highly encouraging that these weakly reflective cells are visible with AO-OCT in line-scan configuration, thus far visualized only with point-scan AO-OCT. Thirdly, line-scan geometry can cause multiple scattering artifacts and cross-talk for imaging structures distal to the highly scattering RPE. That the RPE sub-mosaic is readily visible with line-scan geometry lends support to the notion that multiple scattering artifacts with linear illumination are tolerable, though a thorough analysis of signal roll-off in the retina would help bolster this conclusion. The increased axial resolution allows visualizing a subset of cones with shorter outer segments, suggested to be S-cones [43-45]. These features remained beyond the resolution and sensitivity limits of the lens-based AO line-scan OCT demonstrated previously [16].

Previous studies employing hardware AO and digital aberration correction in OCT have examined cone outer segment elongation in response to a light stimulus, including its application for cone spectral classification [16, 23-26]. Optical phase change is calculated between reflections arising in the ISOS and COST, and averaged over the aperture of the cone to improve sensitivity. Phase noise and eye motion, combined with limited spatial resolution, has restricted these light-evoked observations to the parafoveal cone mosaic. Two deg. temporal eccentricity has been the closest retinal location accessible from the foveal center [25]. Cone spacing and density rise precipitously in the central 1 deg. and pose significant barriers to access. Here, the spatial resolution and speed was sufficient for observing light-evoked responses at eccentricities as close as 0.3 deg. from the foveal center. This enabled the first demonstrations of reduced S-cone density in the foveolar, thus far observed only in histology [45]. Only a single long wavelength stimulus (660 nm) was used here, where one may expect that a dysfunctional cone will exhibit a negligible light-evoked response similar to an S-cone. In Fig. 8c, however, only one such cone with a negligible response was observed while all others (n = 227) exhibited a non-zero response, consistent with their categorization into either L- or M-cones.

In addition to favorable eye motion compensation, the high-speed cross-sectional imaging offered by line-scan OCT enables the observation of fast retinal events [23, 51]. Here, examples of the latter are not shown. Given that the detection arm has the same 2D sensor as our previous reports, the same acquisition speed up to 16 kHz B-scan is possible with this instrument. Still faster burst-mode cameras can provide higher B-scan rates but are more expensive. The benefit for eye motion compensation is evident in revealing weakly reflecting structures and those that benefit from organelle motility, such as the RGC and RPE sub-mosaics. The RGC sub-mosaic here was visualized by averaging 140 - 200 volumes, that is in good agreement with point-scan AO-OCT where 100 - 160 volumes were averaged [8]. Incorporating real-time eye-tracking to stabilize the line-scan OCT probe in real-time on the retina, as has been shown for point-scan OCT [52], will alleviate the deleterious effect of eye motion for phase sensitive applications such as optoretinography and angiography.

In the optical design introduced here, we did not evaluate the extent to which the subject’s own native spherocylindrical prescription degrades theoretical imaging performance. While the optical design remained diffraction limited for a 2.2° field of view, still larger fields of view were not adequately assessed and may eventually require sophisticated optical design engineering to realize. These features will be useful to generalize the feasibility of imaging over a wide population of human subjects with variable refractive error, eye motion and diseased states. For parallel OCT such as in line-scan or full-field configurations, multiple scattering and cross-talk degrade signal quality near and distal to the RPE. Recently, using fast deformable membranes in FF-OCT, significant suppression of cross-talk was shown enabling the visualization of the choroid [53] – a technique that has promise for line-scan implementations as well.

Application of AO-OCT to high-resolution imaging of retinal structure and function in patients with eye disease is rapidly evolving [54-56] and will fuel a surge in our understanding of underlying disease mechanisms on a cellular scale. The common notion that the function of cells degrades earlier than does the structure can now be confirmed with optoretinography. Line-scan AO-OCT is unique in its ability to have access to a wide range of spatiotemporal resolution, wherein both high spatial resolution structure (rods, foveal cones) and high temporal resolution function (ORG early response [23, 51]) can be interrogated in the same platform. Here, with reflective mirror-based telescopes, and theoretically diffraction limited performance, sufficient lateral resolution and sensitivity were obtained to visualize foveal cones, rods, RGCs, and macrophages thus paralleling the imaging performance of point-scan AO-OCT. In addition, high-speed, parallel phase-resolved acquisition enabled classifying the cone spectral types near the foveola, where the reduced density of S-cones observed from prior histology was readily apparent in a living human eye *in vivo*. Given the challenges typically associated with optical accessibility in the living human fovea, the feasibility for high resolution imaging of retinal structure and function demonstrated here holds significant promise for basic and translational applications.

## Acknowledgement and Disclosures

Burroughs Wellcome Fund (Careers at the Scientific Interfaces Award); M.J.Murdock Charitable Trust; Foundation Fighting Blindness; Research to Prevent Blindness (Career Development Award, Unrestriced grant to UW Ophthalmology); National Eye Institute EY027941, EY029710, P30EY001730, U01EY025501).

VPP and RS have a commercial interest in a US patent describing the technology for the line-scan OCT for optoretinography

## References

1. D. Huang, E. A. Swanson, C. P. Lin, J. S. Schuman, W. G. Stinson, W. Chang, M. R. Hee, T. Flotte, K. Gregory, C. A. Puliafito, and et al., “Optical coherence tomography,” Science 254, 1178–1181 (1991).

2. M. Pircher and R. J. Zawadzki, “Review of adaptive optics OCT (AO-OCT): principles and applications for retinal imaging [Invited],” Biomed Opt Express 8, 2536–2562 (2017).

3. D. Miller, J. Qu, R. Jonnal, and K. Thorn, Coherence gating and adaptive optics in the eye, Biomedical Optics (SPIE, 2003), Vol. 4956.

4. B. Hermann, E. J. Fernández, A. Unterhuber, H. Sattmann, A. F. Fercher, W. Drexler, P. M. Prieto, and P. Artal, “Adaptive-optics ultrahigh-resolution optical coherence tomography,” Opt. Lett. 29, 2142–2144 (2004).

5. R. J. Zawadzki, S. M. Jones, S. S. Olivier, M. Zhao, B. A. Bower, J. A. Izatt, S. Choi, S. Laut, and J. S. Werner, “Adaptive-optics optical coherence tomography for high-resolution and high-speed 3D retinal in vivo imaging,” Opt. Express 13, 8532–8546 (2005).

6. Y. Zhang, J. Rha, R. S. Jonnal, and D. T. Miller, “Adaptive optics parallel spectral domain optical coherence tomography for imaging the living retina,” Opt. Express 13, 4792–4811 (2005).

7. D. X. Hammer, A. Agrawal, R. Villanueva, O. Saeedi, and Z. Liu, “Label-free adaptive optics imaging of human retinal macrophage distribution and dynamics,” Proceedings of the National Academy of Sciences 117, 30661–30669 (2020).

8. Z. Liu, K. Kurokawa, F. Zhang, J. J. Lee, and D. T. Miller, “Imaging and quantifying ganglion cells and other transparent neurons in the living human retina,” Proceedings of the National Academy of Sciences of the United States of America 114, 12803–12808 (2017).

9. Z. Liu, J. Tam, O. Saeedi, and D. X. Hammer, “Trans-retinal cellular imaging with multimodal adaptive optics,” Biomed. Opt. Express 9, 4246–4262 (2018).

10. Y. Jian, S. Lee, M. J. Ju, M. Heisler, W. Ding, R. J. Zawadzki, S. Bonora, and M. V. Sarunic, “Lens-based wavefront sensorless adaptive optics swept source OCT,” Scientific reports 6, 27620 (2016).

11. O. P. Kocaoglu, T. L. Turner, Z. Liu, and D. T. Miller, “Adaptive optics optical coherence tomography at 1 MHz,” Biomed. Opt. Express 5, 4186–4200 (2014).

12. D. Merino, C. Dainty, A. Bradu, and A. G. Podoleanu, “Adaptive optics enhanced simultaneous en-face optical coherence tomography and scanning laser ophthalmoscopy,” Opt. Express 14, 3345–3353 (2006).

13. M. Pircher, R. J. Zawadzki, J. W. Evans, J. S. Werner, and C. K. Hitzenberger, “Simultaneous imaging of human cone mosaic with adaptive optics enhanced scanning laser ophthalmoscopy and high-speed transversal scanning optical coherence tomography,” Opt. Lett. 33, 22–24 (2008).

14. R. J. Zawadzki, B. Cense, Y. Zhang, S. S. Choi, D. T. Miller, and J. S. Werner, “Ultrahigh-resolution optical coherence tomography with monochromatic and chromatic aberration correction,” Opt. Express 16, 8126–8143 (2008).

15. C. Torti, B. Považay, B. Hofer, A. Unterhuber, J. Carroll, P. K. Ahnelt, and W. Drexler, “Adaptive optics optical coherence tomography at 120,000 depth scans/s for non-invasive cellular phenotyping of the living human retina,” Opt. Express 17, 19382–19400 (2009).

16. V. P. Pandiyan, X. Jiang, A. Maloney-Bertelli, J. A. Kuchenbecker, U. Sharma, and R. Sabesan, “High-speed adaptive optics line-scan OCT for cellular-resolution optoretinography,” Biomed. Opt. Express 11, 5274–5296 (2020).

17. S.-H. Lee, J. S. Werner, and R. J. Zawadzki, “Improved visualization of outer retinal morphology with aberration cancelling reflective optical design for adaptive optics - optical coherence tomography,” Biomed. Opt. Express 4, 2508–2517 (2013).

18. F. Felberer, J.-S. Kroisamer, B. Baumann, S. Zotter, U. Schmidt-Erfurth, C. K. Hitzenberger, and M. Pircher, “Adaptive optics SLO/OCT for 3D imaging of human photoreceptors in vivo,” Biomed. Opt. Express 5, 439–456 (2014).

19. Z. Liu, O. P. Kocaoglu, and D. T. Miller, “3D Imaging of Retinal Pigment Epithelial Cells in the Living Human Retina,” Investigative Ophthalmology & Visual Science 57, OCT533–OCT543 (2016).

20. M. F. Shirazi, E. Brunner, M. Laslandes, A. Pollreisz, C. K. Hitzenberger, and M. Pircher, “Visualizing human photoreceptor and retinal pigment epithelium cell mosaics in a single volume scan over an extended field of view with adaptive optics optical coherence tomography,” Biomed. Opt. Express 11, 4520–4535 (2020).

21. K. Kurokawa, Z. Liu, and D. T. Miller, “Adaptive optics optical coherence tomography angiography for morphometric analysis of choriocapillaris [Invited],” Biomed Opt Express 8, 1803–1822 (2017).

22. A. Mehdi, V. Denise, V. V. Kari, S. W. John, J. Z. Robert, and S. J. Ravi, “Optoretinogram: optical measurement of human cone and rod photoreceptor responses to light,” Opt. Lett. 45, p4658--4661 (2020).

23. V. P. Pandiyan, A. Maloney-Bertelli, J. A. Kuchenbecker, K. C. Boyle, T. Ling, Z. C. Chen, B. H. Park, A. Roorda, D. Palanker, and R. Sabesan, “The optoretinogram reveals the primary steps of phototransduction in the living human eye,” Science advances 6, eabc1124 (2020).

24. M. Azimipour, J. V. Migacz, R. J. Zawadzki, J. S. Werner, and R. S. Jonnal, “Functional retinal imaging using adaptive optics swept-source OCT at 1.6 MHz,” Optica 6, 300–303 (2019).

25. F. Zhang, K. Kurokawa, A. Lassoued, J. A. Crowell, and D. T. Miller, “Cone photoreceptor classification in the living human eye from photostimulation-induced phase dynamics,” Proceedings of the National Academy of Sciences of the United States of America 116, 7951–7956 (2019).

26. D. Hillmann, H. Spahr, C. Pfäffle, H. Sudkamp, G. Franke, and G. Hüttmann, “In vivo optical imaging of physiological responses to photostimulation in human photoreceptors,” Proceedings of the National Academy of Sciences 113, 13138–13143 (2016).

27. C. Pfäffle, H. Spahr, L. Kutzner, S. Burhan, F. Hilge, Y. Miura, G. Hüttmann, and D. Hillmann, “Simultaneous functional imaging of neuronal and photoreceptor layers in living human retina,” Opt. Lett. 44, 5671–5674 (2019).

28. R. F. Cooper, D. H. Brainard, and J. I. W. Morgan, “Optoretinography of individual human cone photoreceptors,” Opt. Express 28, 39326 (2020).

29. R. F. Cooper, W. S. Tuten, A. Dubra, D. H. Brainard, and J. I. W. Morgan, “Non-invasive assessment of human cone photoreceptor function,” Biomed. Opt. Express 8, 5098–5112 (2017).

30. R. S. Jonnal, J. Rha, Y. Zhang, B. Cense, W. Gao, and D. T. Miller, “In vivo functional imaging of human cone photoreceptors,” Opt. Express 15, 16141–16160 (2007).

31. R. S. Jonnal, J. R. Besecker, J. C. Derby, O. P. Kocaoglu, B. Cense, W. Gao, Q. Wang, and D. T. Miller, “Imaging outer segment renewal in living human cone photoreceptors,” Opt Express 18, 5257–5270 (2010).

32. O. P. Kocaoglu, Z. Liu, F. Zhang, K. Kurokawa, R. S. Jonnal, and D. T. Miller, “Photoreceptor disc shedding in the living human eye,” Biomed. Opt. Express 7, 4554–4568 (2016).

33. K. Grieve and A. Roorda, “Intrinsic signals from human cone photoreceptors,” Investigative ophthalmology & visual science 49, 713–719 (2008).

34. P. Bedggood and A. Metha, “Variability in bleach kinetics and amount of photopigment between individual foveal cones,” Invest Ophthalmol Vis Sci 53, 3673–3681 (2012).

35. P. Bedggood and A. Metha, “Optical imaging of human cone photoreceptors directly following the capture of light,” PloS one 8(2013).

36. R. Sabesan, H. Hofer, and A. Roorda, “Characterizing the Human Cone Photoreceptor Mosaic via Dynamic Photopigment Densitometry,” PLoS One 10, e0144891 (2015).

37. A. Dubra and Y. Sulai, “Reflective afocal broadband adaptive optics scanning ophthalmoscope,” Biomedical optics express 2, 1757–1768 (2011).

38. A. Gomez-Vieyra, A. Dubra, D. Malacara-Hernandez, and D. R. Williams, “First-order design of off-axis reflective ophthalmic adaptive optics systems using afocal telescopes,” Opt Express 17, 18906–18919 (2009).

39. M. Laslandes, M. Salas, C. K. Hitzenberger, and M. Pircher, “Increasing the field of view of adaptive optics scanning laser ophthalmoscopy,” Biomed. Opt. Express 8, 4811–4826 (2017).

40. M. Jensen, I. B. Gonzalo, R. D. Engelsholm, M. Maria, N. M. Israelsen, A. Podoleanu, and O. Bang, “Noise of supercontinuum sources in spectral domain optical coherence tomography,” JOSA B 36, A154–A160 (2019).

41. A. Dubra and Y. Sulai, “Reflective afocal broadband adaptive optics scanning ophthalmoscope,” Biomed. Opt. Express 2, 1757–1768 (2011).

42. C. W. Oyster, “The human eye,” Sunderland, MA: Sinauer (1999).

43. D. Valente, M. Azimipour, R. J. Zawadzki, J. S. Werner, and R. Jonnal, “Investigating the morphology of possible S-cones using adaptive optics functional OCT,” Investigative Ophthalmology & Visual Science 60, 4595–4595 (2019).

44. D. T. Miller and K. Kurokawa, “Cellular-scale imaging of transparent retinal structures and processes using adaptive optics optical coherence tomography,” Annual Review of Vision Science 6, 115–148 (2020).

45. C. A. Curcio, K. A. Allen, K. R. Sloan, C. L. Lerea, J. B. Hurley, I. B. Klock, and A. H. Milam, “Distribution and morphology of human cone photoreceptors stained with anti-blue opsin,” Journal of Comparative Neurology 312, 610–624 (1991).

46. Z. Liu, O. P. Kocaoglu, T. L. Turner, and D. T. Miller, “Modal content of living human cone photoreceptors,” Biomed. Opt. Express 6, 3378–3404 (2015).

47. Z. Liu, K. Kurokawa, D. X. Hammer, and D. T. Miller, “In vivo measurement of organelle motility in human retinal pigment epithelial cells,” Biomed. Opt. Express 10, 4142–4158 (2019).

48. T. Liu, H. Jung, J. Liu, M. Droettboom, and J. Tam, “Noninvasive near infrared autofluorescence imaging of retinal pigment epithelial cells in the human retina using adaptive optics,” Biomed. Opt. Express 8, 4348–4360 (2017).

49. Z. Liu, O. P. Kocaoglu, and D. T. Miller, “In-the-plane design of an off-axis ophthalmic adaptive optics system using toroidal mirrors,” Biomed. Opt. Express 4, 3007–3030 (2013).

50. K. Kurokawa, J. A. Crowell, N. Do, J. J. Lee, and D. T. Miller, “Multi-reference global registration of individual A-lines in adaptive optics optical coherence tomography retinal images,” Journal of Biomedical Optics 26, 016001 (2021).

51. K. C. Boyle, Z. C. Chen, T. Ling, V. P. Pandiyan, J. Kuchenbecker, R. Sabesan, and D. Palanker, “Mechanisms of Light-Induced Deformations in Photoreceptors,” Biophys J 119, 1481–1488 (2020).

52. B. Braaf, K. V. Vienola, C. K. Sheehy, Q. Yang, K. A. Vermeer, P. Tiruveedhula, D. W. Arathorn, A. Roorda, and J. F. de Boer, “Real-time eye motion correction in phase-resolved OCT angiography with tracking SLO,” Biomed Opt Express 4, 51–65 (2013).

53. E. Auksorius, D. Borycki, and M. Wojtkowski, “Crosstalk-free volumetric in vivo imaging of a human retina with Fourier-domain full-field optical coherence tomography,” Biomed. Opt. Express 10, 6390–6407 (2019).

54. Z. Liu, O. Saeedi, F. Zhang, R. Villanueva, S. Asanad, A. Agrawal, and D. X. Hammer, “Quantification of retinal ganglion cell morphology in human glaucomatous eyes,” Investigative ophthalmology & visual science 62, 34–34 (2021).

55. A. Lassoued, F. Zhang, K. Kurokawa, Y. Liu, J. A. Crowell, and D. T. Miller, “Measuring dysfunction of cone photoreceptors in retinitis pigmentosa with phase-sensitive AO-OCT,” in Ophthalmic Technologies XXX, (International Society for Optics and Photonics, 2020), 1121815.

56. F. Zhang, K. Kurokawa, M. T. Bernucci, H. W. Jung, A. Lassoued, J. A. Crowell, J. Neitz, M. Neitz, and D. T. Miller, “Revealing How Color Vision Phenotype and Genotype Manifest in Individual Cone Cells,” Investigative Ophthalmology & Visual Science 62, 8–8 (2021).

